# Clinal and seasonal change are correlated in *Drosophila melanogaster* natural populations

**DOI:** 10.1101/2020.03.19.999011

**Authors:** Murillo F. Rodrigues, Maria D. Vibranovski, Rodrigo Cogni

**Affiliations:** Department of Genetics and Evolutionary Biology, Institute of Biosciences, University of Sao Paulo, Sao Paulo, SP, Brazil; Institute of Ecology and Evolution, University of Oregon, Eugene, OR, USA; Department of Ecology, Institute of Biosciences, University of Sao Paulo, Sao Paulo, SP, Brazil

## Abstract

Spatial and seasonal variation in the environment are ubiquitous. Environmental heterogeneity can affect natural populations and lead to covariation between environment and allele frequencies. *Drosophila melanogaster* is known to harbor polymorphisms that change both with latitude and seasons. Identifying the role of selection in driving these changes is not trivial, because non-adaptive processes can cause similar patterns. Given the environment changes in similar ways across seasons and along the latitudinal gradient, one promising approach may be to look for parallelism between clinal and seasonal change. Here, we test whether there is a genome-wide correlation between clinal and seasonal change, and whether the pattern is consistent with selection. Allele frequency estimates were obtained from pooled samples from seven different locations along the east coast of the US, and across seasons within Pennsylvania. We show that there is a genome-wide correlation between clinal and seasonal variation, which cannot be explained by linked selection alone. This pattern is stronger in genomic regions with higher functional content, consistent with natural selection. We derive a way to biologically interpret these correlations and show that around 3.7% of the common, autosomal variants could be under parallel seasonal and spatial selection. Our results highlight the contribution of natural selection in driving fluctuations in allele frequencies in natural fly populations and point to a shared genomic basis to climate adaptation which happens over space and time in *D. melanogaster*.

## Introduction

Species occur in environments that vary both in space and time (Ewing 1979; Cardini et al. 2007; Dionne et al. 2007; Hancock et al. 2008; Zuther et al. 2012; Campitelli and Stinchcombe 2013; Kooyers et al. 2015). Populations may adapt to the local conditions of the environment in which they occur, resulting in covariation between traits and space (Endler 1977; Barton 1983; Barton 1999; Kawecki and Ebert 2004). Similarly, predictable changes in the environment through time can lead covariation between relevant traits and time (Levene, 1953; Ewing 1979). Although correlations between environment and traits (either in time or space) are indicative of selection, these patterns can be produced by non-adaptive processes such as migration, isolation by distance and range expansion (Wright 1943; Vasemägi 2006; Excoffier et al. 2009; Duchen et al. 2013; Bergland et al. 2016). It is not trivial to identify the role of selection in diversifying traits, but a promising approach might be to jointly model changes across space and time.

*Drosophila melanogaster* is a uniquely suited to study both spatial and temporal adaptation. These sub-Saharan flies recently invaded most of the world (David and Capy 1988), and adaptations at the phenotypic and genotypic levels evolved in response to the colonization of new habitats (Mettler et al. 1977; Knibb 1982; Oakeshott et al. 1982; David and Capy 1988; Schmidt et al. 2000; de Jong and Bochdanovits 2003; Sezgin et al. 2004). Many traits, polymorphisms and inversions were observed to covary with latitude (also called clinal) in natural fly populations (Hoffmann et al. 2002; Hoffmann and Weeks 2007; Turner et al. 2008; Paaby et al. 2010; Yukilevich et al. 2010; Reinhardt et al. 2014; Schrider et al. 2016). For instance, flies from colder environments are darker (David et al. 1985), bigger (Arthur et al. 2008) and show higher incidence of reproductive diapause than flies from lower latitudes (Schmidt et al. 2005).

In higher latitudes, fly populations started to experience dramatic cyclical changes in the environment through seasons. Given these flies have multiple generations per year, differential fitness across seasons could theoretically lead to temporal adaptations (Levene 1953; Ewing 1979). Traits that favor rapid reproduction in the summer can be particularly different to those which favor endurance in the winter (Behrman et al. 2015). Concordant with this hypothesis, chromosomal inversions in *D. pseudoobscura* were observed to cycle with seasons (Dobzhansky 1943). In *D. melanogaster*, flies collected in the spring are more tolerant to stress (Behrman et al. 2015), show higher diapause inducibility (Schmidt and Conde 2006), have increased immune function (Behrman et al. 2018) and have different cuticular hydrocarbon profiles than those collected in the fall (Rajpurohit et al. 2017). Genome-wide analyses have identified polymorphisms and inversions that oscillate in seasonal timescales in several localities in the United States and Europe (Bergland et al. 2014; Kapun et al. 2016; Machado et al. 2021). However, a recent analysis suggested seasonal fluctuations in allele frequencies seems small and temporal structure independent of seasons may be more important in this system (Buffalo and Coop 2019).

It is challenging to characterize the role of selection in producing spatial or seasonal change in allele frequencies. At the spatial scale, the axis of demography and environmental heterogeneity are confounded in this system (i.e., migration and environment are structured along the south-north axis) (Caracristi and Schlotterer 2003; Yukilevich and True 2008; Duchen et al. 2013; Kao et al. 2015). At the seasonal scale, the magnitude of allele frequency change with seasons is expected to be rather small, and stochastic environmental events not aligned with seasons complicate inferences even further (Machado et al. 2021). However, we can gain power by jointly modelling latitudinal and seasonal changes (Cogni et al. 2015).

Two adaptive mechanisms are expected to induce correlations between clinal and seasonal fluctuations in allele frequencies. First, the environment changes similarly with latitude and through seasons (at least with respect to temperature). Second, the onset of spring changes with latitude, and so seasonal changes in polymorphisms alone could produce clinal variation, a mechanism termed seasonal phase clines (Roff 1980; Rhomberg and Singh 1986). The effects of neutral, demographic processes which can confound interpretation are expected to be much less pronounced because of the short time scale of seasonal processes. Thus, parallel latitudinal and seasonal variation in a trait is strong evidence in favor of natural selection (Bergland et al. 2014; Cogni et al. 2015).

Some empirical studies have found parallelism between clinal and seasonal variation in *D. melanogaster* (Bergland et al. 2014; Cogni et al. 2015; Kapun et al. 2016; Behrman et al. 2018; Machado et al. 2021). The prevalence of reproductive diapause, a phenotype tightly linked to adaptation to cold environments, varies both with latitude and seasons (Schmidt et al. 2005). Cogni et al. (2014) found that a variant in the *couchpotato* gene, which encodes diapause inducibility, also varied predictably with latitude and across seasons: the diapause-inducing allele is positively correlated with latitude and its frequency increases from summer into winter. Cogni et al. (2015) found an association between clinal and seasonal change in central metabolic genes, which are likely important drivers of climatic adaptation. Kapun et al. (2016) found that a few cosmopolitan inversions thought to be involved with climate adaptation also vary in parallel with latitude and through seasons.

Here, we test whether parallel clinal and seasonal variation is pervasive across the *D. melanogaster* genome. It is essential we further our understanding of the genomic basis to climate adaptation in *D. melanogaster*, so that we can identify possible mechanisms which allow adaptation over such short time scales (Wittmann et al. 2017). A parallel and independent study also investigated the relationship between clinal and seasonal change in *D. melanogaster* (Machado et al. 2021). Nevertheless, our work is fundamentally different from previous studies because (i) we use seasonal samples collected over six years, as opposed to at most three years in other studies; (ii) our samples are all from the same location in Pennsylvania, where seasonality is strong and phenotypes are known to cycle seasonally, and where there is little evidence of population substructure or large scale migrations events (Schmidt and Conde 2006; Behrman et al. 2015; Rajpurohit et al. 2017; Behrman et al. 2018); (iii) we analyze how the correlation between clinal and seasonal variation changes across genomic regions which differ in density of functional sites, allowing us to better disentangle demography and selection; and (iv) we dissect the role of linkage disequilibrium in driving these patterns.

## Material and Methods

### Population samples

We analyzed 20 samples from seven locations along the United States east coast, collected by (Bergland et al. 2014) (10 samples), and (Machado et al. 2021) (10 samples) (see Table S1). The samples were based on pools of wild-caught individuals. We decided to not include previously collected samples from Maine because they were collected in the fall, whereas all of our other samples were collected in the spring, and we also did not include the DGRP sample from North Carolina, as it is hard to ascertain when they were obtained (Fabian et al. 2012; Mackay et al. 2012; Bergland et al. 2014). The Linvilla (Pennsylvania) population was sampled extensively from 2009 to 2015 (six spring, seven fall samples), and was therefore used in our analysis of seasonal variation. One of the Pennsylvania samples was exclusively used for the clinal analysis to minimize dependency between our clinal and seasonal sets. We also replicated our clinal analysis using data from four Australian samples (Anderson et al. 2005; Kolaczkowski et al. 2011). All the data used in this project are available on the NCBI Short Read Archive (BioProject accession numbers PRJNA256231, PRJNA308584 and NCBI Sequence Read Archive SRA012285.16).

### Mapping and processing of sequencing data

Raw, paired-end reads were mapped against the FlyBase *D. melanogaster* (r6.15) and *D. simulans* (r2.02) reference genomes (Gramates et al. 2017) using BBSplit from the BBMap suite (https://sourceforge.net/projects/bbmap/; version from February 11, 2019). We removed any reads that preferentially mapped to *D. simulans* to mitigate effects of contamination (the proportion of reads preferentially mapping to *D. simulans* was minimal, never exceeding 3%). Then, reads were remapped to *D. melanogaster* reference genome using bwa (MEM algorithm) version 0.7.15 (Li and Durbin 2010). Files were converted from SAM to BAM format using Picard Tools (http://broadinstitute.github.io/picard). PCR duplicates were marked and removed using Picard Tools and local realignment around indels was performed using GATK version 3.7 (McKenna et al. 2010). Single nucleotide polymorphisms (SNPs) and indels were called using CRISP with default parameters (Bansal et al. 2016).

We applied several filters to ensure that the identified SNPs were not artifacts. SNPs in repetitive regions, identified using the RepeatMasker library for *D. melanogaster* (obtained from http://www.repeatmasker.org), and SNPs within 5bp of polymorphic indels were removed from our analyses. SNPs with mean minor allele frequency in the clinal and seasonal samples less than 5%, with minimum per-population coverage less than 10x (or 4x for the Australian samples) or maximum per-population coverage greater than the 99^th^ quantile were excluded from our analyses. We only considered bi-allelic, autosomal SNPs in our downstream analyses. Functional annotations for the identified SNPs obtained using SNPeff version 4.3o (Cingolani et al. 2012).

### Clinal and seasonal changes in allele frequency

The allele frequencies were calculated by diving the number of reads supporting each allele, divided by the total number of reads. Because pool-seq data contain an additional component of error due to sampling, we did not weight the allele frequencies by total depth at each site; instead we used the effective sample size, or effective number of chromosomes (*N_E_*), as the denominator. This metric can be computed as follows:

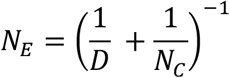

where *N_C_* is the number of chromosomes in the pool and *D* is the read depth at that site (Kolaczkowski et al. 2011; Feder et al. 2012; Bergland et al. 2014).

To assess latitudinal variation, we fitted a binomial linear model of allele frequency against latitude for each site. Similarly, we regressed allele frequency at each site against a season dummy variable (June and July were encoded as Spring, and September, October and November as Fall) and included the year of sampling as a covariate. For either regression, we required the variant to be polymorphic in at least two samples. Further, we computed pairwise F_ST_ for all our samples using the R package poolfstat (Hivert et al., 2018).

We defined clinal and seasonal SNPs using an outlier approach, because we do not have an adequate genome-wide null distribution to compare our estimates. We considered that SNPs were outliers if their regression P-value fell in the bottom 1% (or 5%) of the distribution.

### Correlation between clinal and seasonal variation

Our main goal was to evaluate whether clinal and seasonal change are correlated, pooling information across the thousands of polymorphisms that segregate in natural populations. To do so, we regressed the slopes of the clinal regressions and the slope of the seasonal regressions. The regression line was fit using Huber’s M estimator to improve robustness to outliers. Before fitting the regression, we z-normalized the clinal and seasonal slopes, so the slope of the regression of clinal and seasonal change is actually the same as the correlation.

We also investigated how the correlation between clinal and seasonal change differed across genomic regions. For that, we used a dummy variable with annotations as a covariate. The regions analyzed were exon, intron, 5’ UTR, 3’ UTR, upstream, downstream intergenic and splice. There are some chromosomal inversions segregating in the populations we studied, and they are known to contribute to adaptation (Wright and Dobzhansky 1946; García-Vázquez and Sánchez-Refusta 1988; Kapun et al. 2014). We annotated SNPs surrounding (2Mb) common inversion breakpoints and added inversion status as a covariate in the linear model (Corbett-Detig and Hartl 2012).

To confirm our results are robust to potential model misspecifications, we implemented a permutation test in which we rerun the regressions for each SNP using shuffled season and latitude labels 2,000 times. The same procedure was implemented for most of the statistical tests, except where indicated otherwise.

### Enrichment tests

We tested for enrichment of genic classes using our sets of clinal and seasonal SNPs using Fisher’s exact test for each genic region and statistic. To control for confounders, such as read depth and allele frequency variation, we shuffled the season and latitude labels and reran the generalized regressions. Using the P-values obtained from regressions in which season and latitude labels were shuffled, we defined, for each iteration, lists of top clinal and seasonal SNPs. Then, we calculated the enrichment of each genic class using Fisher’s exact test. To obtain a P-value for an enrichment of a given genic class, we compared the observed odds ratio in the actual dataset to the distribution of odds ratios observed for datasets in which season and latitude labels were shuffled.

### Mitigating the impact of linkage disequilibrium

Selection at one site affects genetic variation at nearby, linked neutral sites (Smith and Haigh 1974). Because we assume that sites are independent in our models, the indirect effects of selection can inflate the magnitude of the patterns we investigated. To test the effect of linkage disequilibrium (LD) in our outlier analyses, we plotted P-values against distance to a top SNP. We then smoothed the scatterplot using cubic splines as implemented in ggplot2 (Wickham 2016). To test the effect of linkage on the relationship between clinal and seasonal variation, we implemented a thinning approach. Sampling one SNP per *L* base pairs one thousand times, we constructed sets of SNPs with minimized dependency, where *L* ranged from 1 to 20kb. For each of these sets for a given *L*, we computed the correlation between clinal and seasonal slopes, and compared the distribution of the thinned regression coefficients to the coefficients we obtained using all SNPs.

All statistical analyses were performed in R 3.5.0 (R Core Team 2018) and can be found at gitlab.com/mufernando/clinal_sea.git.

## Results

We assembled 20 *D. melanogaster* population samples collected from seven localities across multiple years in the east coast of the United States. All of these samples are the result of a collaborative effort of many researchers from a consortium, the DrosRTEC (Bergland et al. 2014; Machado et al. 2021). Seven of our samples span from Florida to Massachusetts and together comprise our clinal set. The seasonal samples were collected in Pennsylvania in the spring (6 samples collected in June or July) and in the fall (6 samples collected in September, October or November). For each sample, a median of 55 individuals (with a range of 33 to 116) was pooled and resequenced to an average 75x coverage (ranging from 17 to 215). We also used four clinal samples from the Australia (Anderson et al. 2005; Kolaczkowski et al. 2011). More details about the samples can be found on Table S1 (also see Machado et al. 2021; Bergland et al. 2014). After all the filtering steps, we identified 798,176 common autosomal SNPs, which were used in our downstream analyses.

### Allele frequency changes with latitude, seasons and years

Latitude explains much of allele frequency variation along the surveyed populations, as there is an excess of low GLM P-value SNPs (Fig. 1A). The mean absolute difference in allele frequency between the ends of the clines is 9.2%. Seasons, on the other hand, explain less of the variation in allele frequency. There is only a minor excess of low GLM P-value SNPs (Fig. 1B) and the mean absolute difference in allele frequency between seasons is 2.6%. We also found that year of sampling is a good predictor of allele frequency change (in Pennsylvania), more so than seasons, given there is a huge excess of low GLM P-values (Fig. 1C).

**Figure 1.**
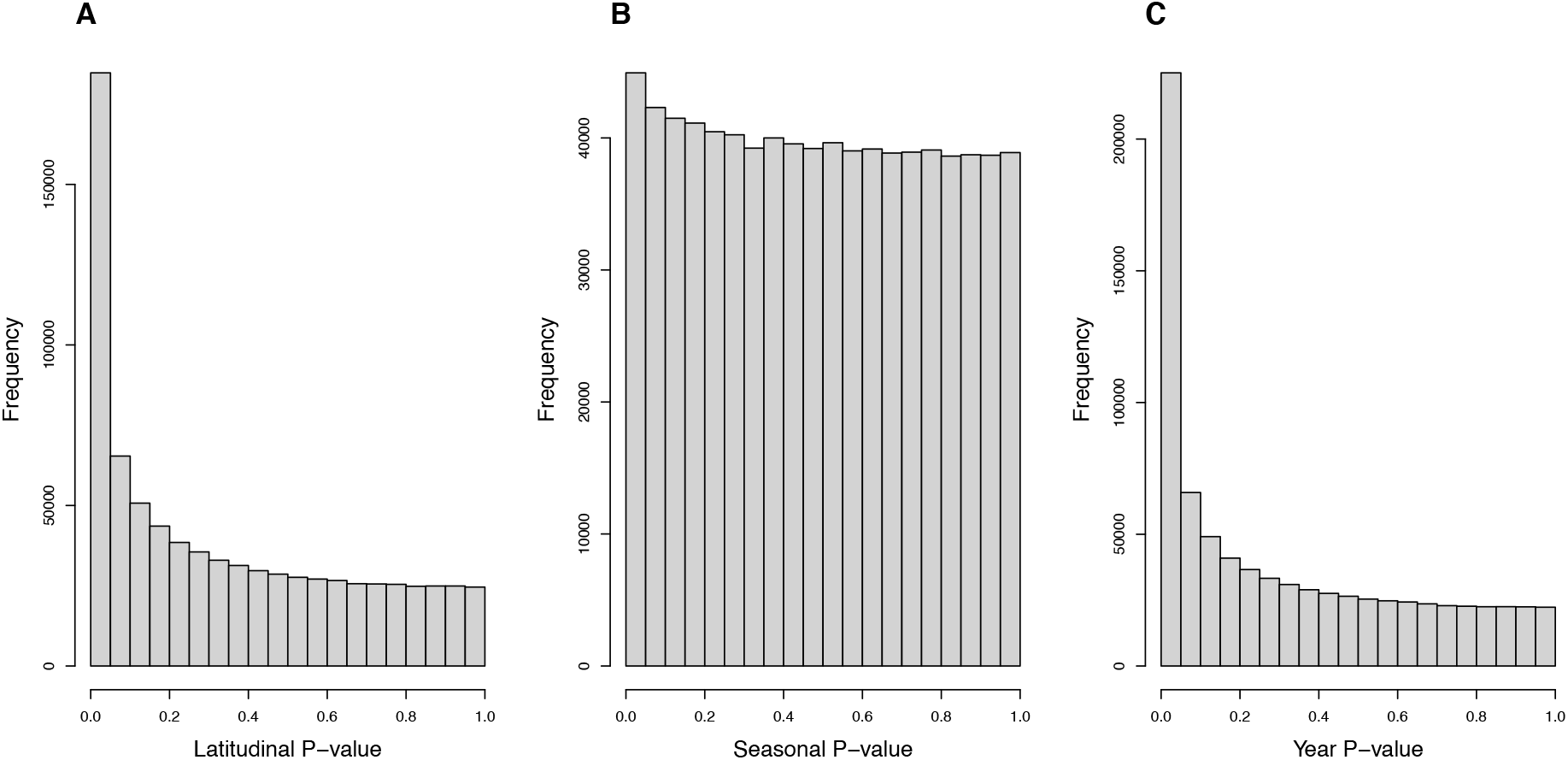
Distribution of P-values from the generalized linear models of allele frequency and latitude, and allele frequency and seasons/years.

Our generalized linear models do not account for dependency between samples, which can be a problem when regressing allele frequency on seasons. To investigate whether this could be an issue, we performed Durbin-Watson tests for autocorrelation in the residuals of the seasonal regressions using Julian days as the time variable. There is no excess of low P-values (Fig. S1), and the season P-values are not correlated with Durbin-Watson test P-value (*P* = 0.77). This indicates that the assumption of independency is being met for most variants, and that autocorrelation is not artificially creating patterns of seasonality in allele frequency.

Given we do not have enough information to build an appropriate null distribution to calibrate our P-values, we sought to demonstrate that top significant clinal and seasonal SNPs are enriched for functional variants, which are more likely to contribute to adaptation. Latitudinal SNPs are more likely to be in exonic and UTR 3’ regions (Fig. 2A), whereas seasonal SNPs are enriched for exonic, UTR 3’ and UTR 5’ regions (Fig. 2B). Further, top latitudinal and seasonal SNPs seem to be underrepresented within upstream and downstream regions. Similar enrichment patterns have been observed for both top clinal and seasonal (Kolaczowski et al. 2011; Fabian et al. 2012,; Bergland et al. 2014; Machado et al. 2016, 2021). Using a 5% cutoff, our enrichment results are largely replicated (Fig. S2).

**Figure 2.**
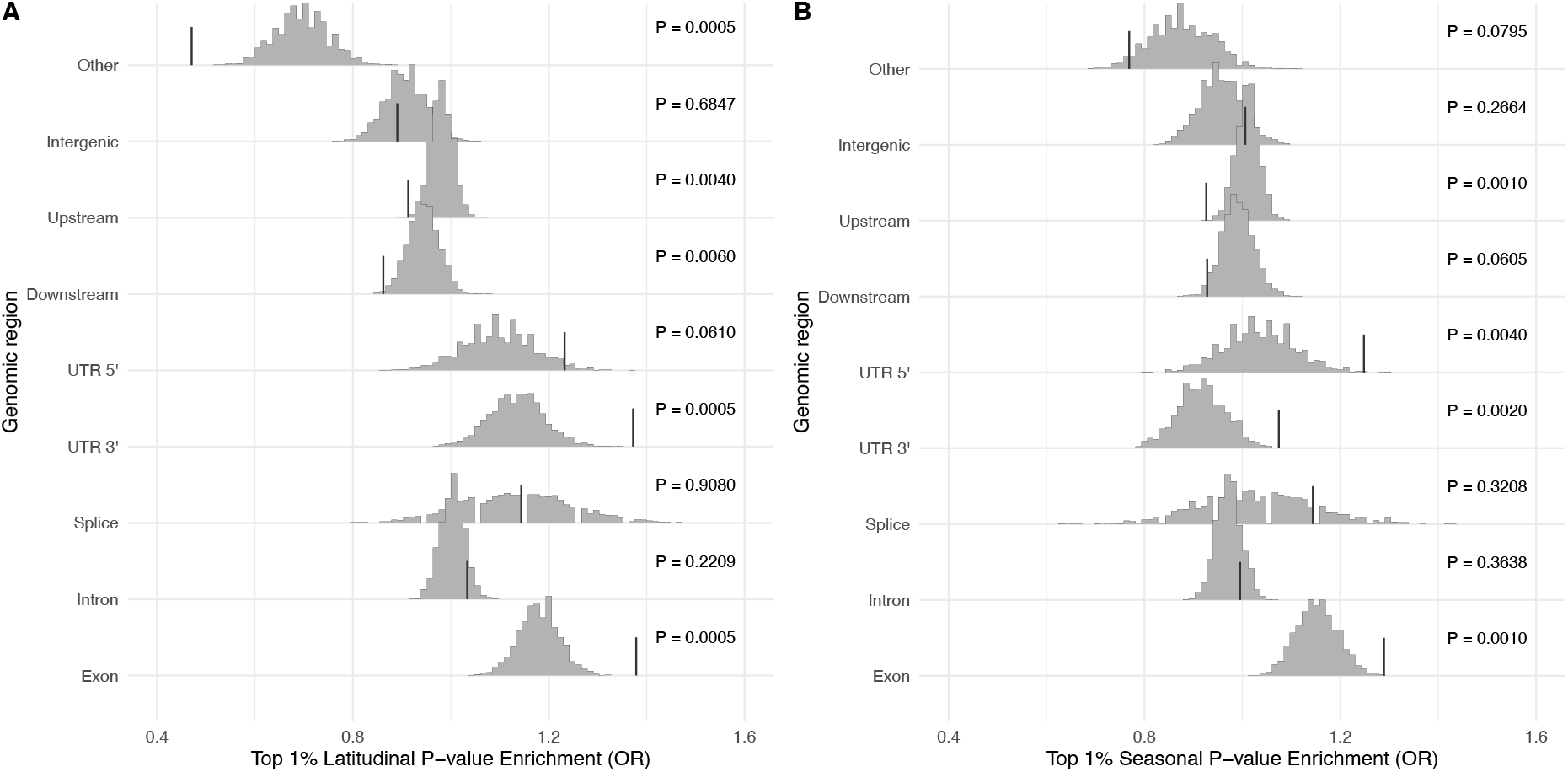
Top SNPs are enriched for functionally relevant classes. Enrichment of top 1% SNPs in each genic class for A) latitudinal P-value and B) seasonal P-value. The histograms show the distribution of odds ratios when latitude and season labels were permuted, and the vertical bars show the observed odds ratios.

### Clinal variation is related to seasonal variation

A clinal pattern can arise solely as a result of demographic processes, such as isolation by distance and admixture (Duchen et al. 2013; Kao et al. 2015; Bergland et al. 2016). Although seasonality is less affected by such processes, seasonal change is less pronounced and more subject to stochastic changes in the environment, making it harder to detect seasonal change with precision. Here, we integrate both clinal and seasonal change estimates across a large number of SNPs in the genome. We expect the overall pattern that emerges to be informative of the relative role of natural selection, because selection is a plausible process to produce a pattern of clinal variation mirroring seasonal variation (Cogni et al. 2015).

We found a significant negative correlation between clinal and seasonal regression coefficients (Fig. 3A, Table S2). The correlation is strongest for SNPs within exons, and the weakest for unclassified SNPs and those within intergenic regions (Fig. 3A; Table S3). Nonetheless, the correlation is different than zero for all classes (except for the unclassified), what would be consistent either with pervasive linked selection or widespread distribution of variants that are important for adaptation, even within non-coding regions. Qualitatively similar results were replicated using a different minor allele frequency cutoff and using samples from Maine, which were obtained the summer – in contrast to all the other clinal populations that were sampled in the fall (Fig. S3B-C).

**Figure 3.**
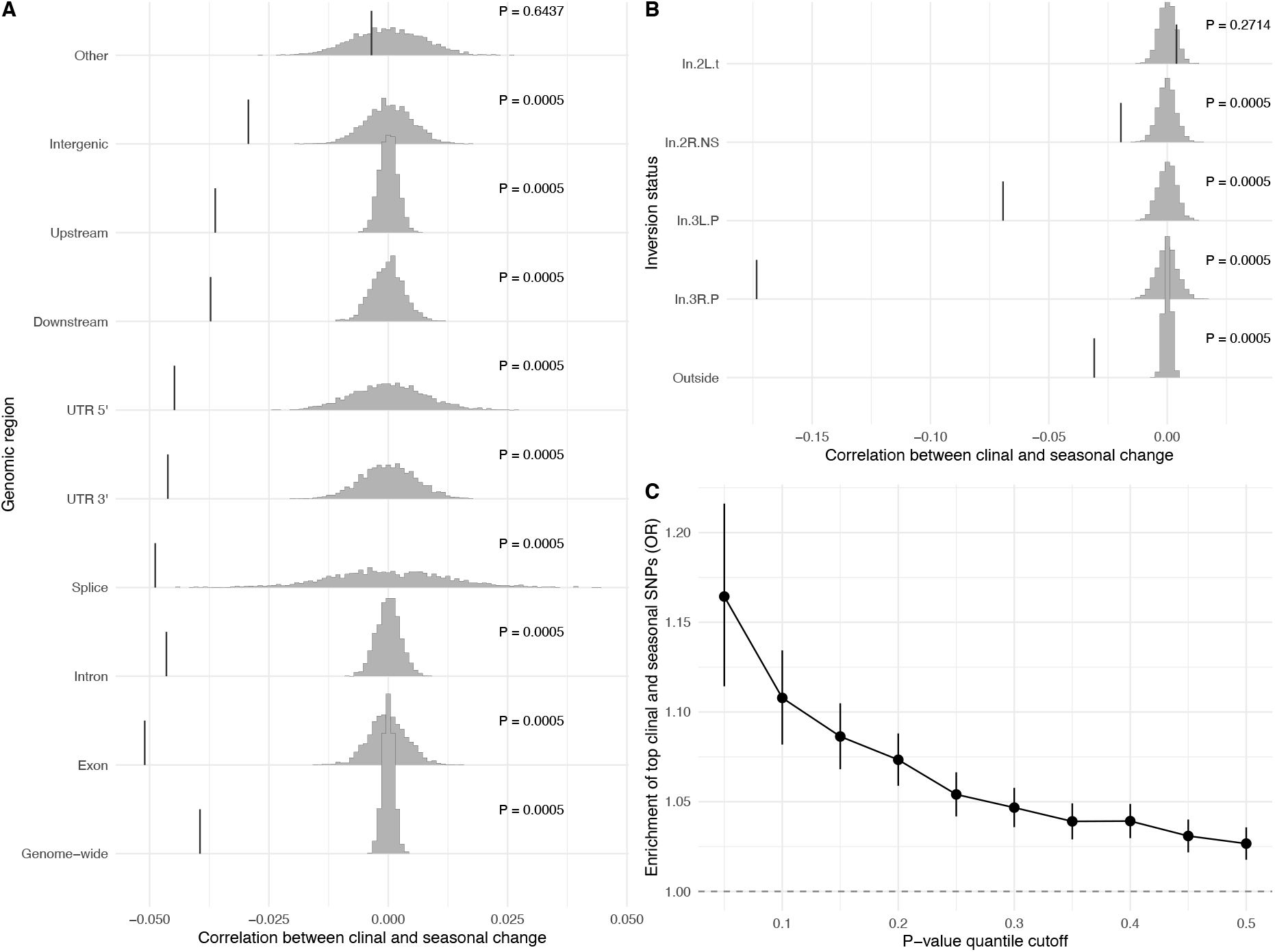
Clinal change parallels seasonal change. Correlation between clinal change and seasonal change for each genic class (A) and by inversion breakpoint (B). (C) Association between top latitudinal and seasonal variants for different P-value cutoffs. Gray histograms are the null distribution of the correlation (after permuting the latitude and season labels) and vertical bars represent the observed correlation.

Given that previous studies have demonstrated the importance of cosmopolitan inversions in climatic adaptation (e.g., Kapun et al. 2016), we looked at the correlation between clinal and seasonal change near common cosmopolitan inversions breakpoints. We found that the correlation between clinal and seasonal change is strongest near the breakpoints of inversions In(2R)NS, In(3R)P and In(3L)P (Fig. 3B, Table S4). Nevertheless, the pattern is still strong outside these regions, indicating our main results are not purely driven by frequency changes of inversions.

Another way of testing for parallelism between clinal and seasonal change is by testing if clinal SNPs are more likely to be seasonal (and vice-versa). We observed that clinal SNPs are enriched for seasonal SNPs (Fig. 3C). The enrichment increases with more stringent lower P-value quantile cut-offs, as we would expect if even strictly non-significant variants were informative of the role of selection.

We also confirmed our main finding, that clinal and seasonal change are correlated using clinal samples from Australia. To measure clinal change in Australia, we only used four low coverage samples in Australia, two for each of low and high latitude locations. There is a negative correlation between clinal variation in Australia and seasonal variation in Pennsylvania that, although minor, is significant (Fig. S3C).

### The effects of linkage disequilibrium on clinal and seasonal variation

Variation at one site is linked to variation at other sites, and selection will increase this dependency (Smith and Haigh 1974). First, we assessed if latitudinal and seasonal P-values, were dependent on how distant a SNP was from our top 1% SNPs. We show that both statistics are dependent on distance from the outlier SNPs (Fig. 4A, B), but the effect virtually disappears after 5kb.

**Figure 4.**
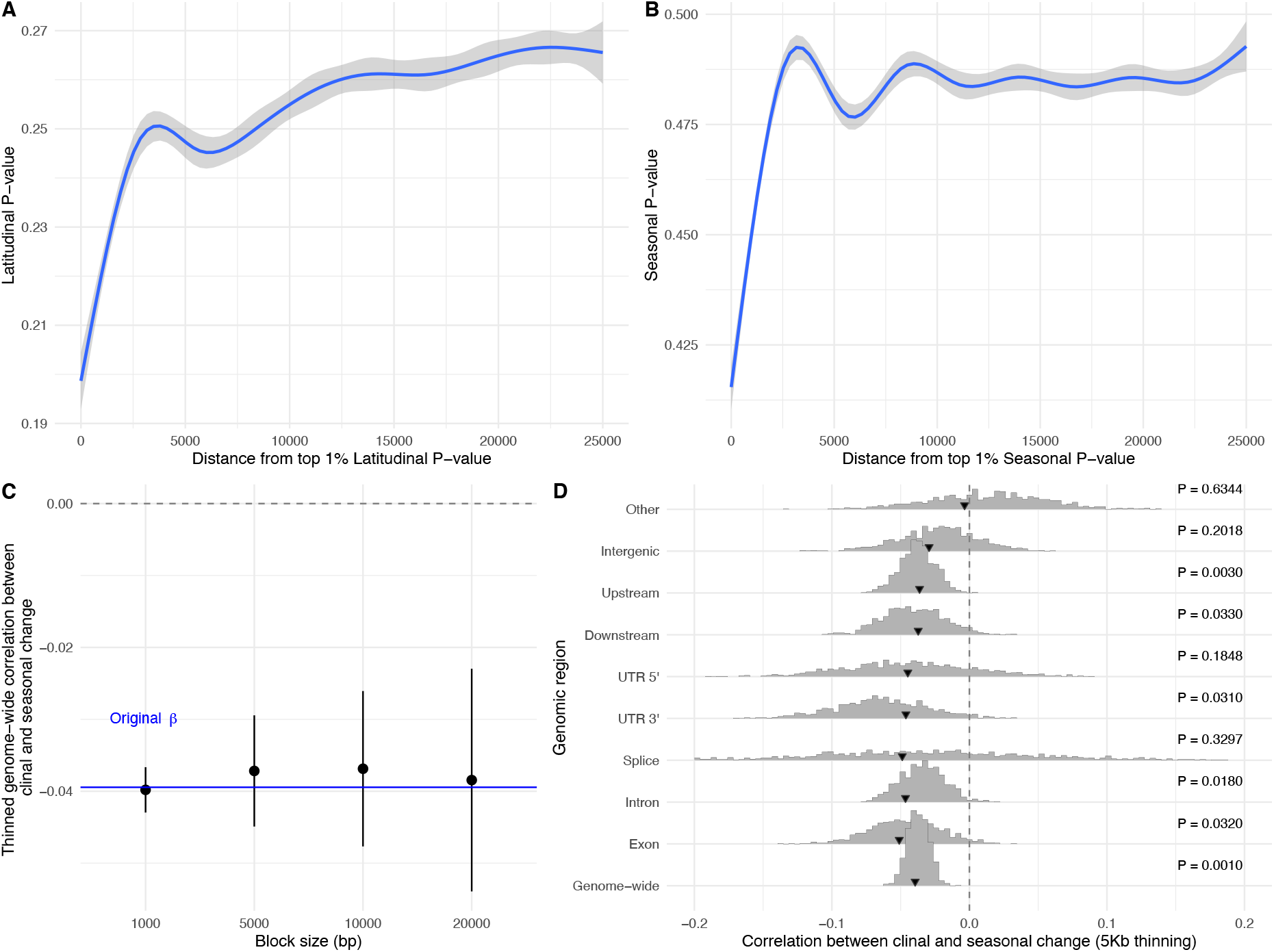
Effects of linkage disequilibrium. The mean (A) latitudinal P-value and (B) seasonal P-value depend on distance to the respective top 1% outliers. The correlation between clinal and seasonal variation is affected by dependency among SNPs. C) the correlation between clinal and seasonal variation changes with the size of the thinning window. D) comparison among original estimates (arrows) and values obtained after thinning using a window size of 5kb (histogram). Histograms show the distributions across sampled thinned datasets, and the black arrows point to the original estimates.

We assessed the impact of linkage on the correlation between clinal and seasonal change by implementing a thinning approach. First, we tested how the genome-wide regression estimate varied with changing window sizes. The effect of non-independency of variants on the correlation is rather small (Fig. 4C) and the genome-wide correlation remains significantly different from zero (P= 0.001; Fig. 4D). The thinning did not significantly reduce the signal for many regions, but the strength of the signal within splicing, UTR 5’, upstream, downstream and intergenic regions decreased and did not remain significantly different from zero (Fig. 4D). It seems that most of the signal coming from those regions are due to the linked effects of selection.

### Biological interpretation of the correlation between clinal and seasonal change

Although the negative correlation between clinal and seasonal change indicates a role for selection, it is unclear how strongly parallel selection would need to be to generate this correlation. Intuitively, we expect the correlation to be rather small, as the majority of the variants are likely not under parallel selection. Below, we derive how to get a rough estimate of the number of SNPs under parallel selection from the observed correlation.

Suppose that for a proportion *p* of the SNPs the clinal and seasonal regression coefficients are correlated due to parallel selection, whereas for the remainder of the SNPs they are independent. What genome-wide correlation would we expect? To find this, we can write (*Z*_1_, *Z*_2_) for the two z-normalized regression coefficients of a randomly chosen SNP. SNPs can be either under parallel selection or not, so we define (*X*_1_, *X*_2_) and (*Y*_1_, *Y*_2_) as random draws from the z-normalized regression coefficients for SNPs under parallel selection and not, respectively. Then, (*Z*_1_, *Z*_2_) = (*X*_1_, *X*_2_) with probability *p*, and (*Z*_1_, *Z*_2_) = (*Y*_1_, *Y*_2_) otherwise. To find the cor(*Z*_1_, *Z*_2_), note that since *Z*_1_ and *Z*_2_ have mean 0 and standard deviation 1, cor(*Z*_1_, *Z*_2_) = *E*[*Z*_1_*Z*_2_] (and similarly for *X* and *Y*). So, cor(*Z*_1_, *Z*_2_) = *p* cor(*X*_1_, *X*_2_) + (1 – *p*) cor(*Y*_1_, *Y*_2_) using the law of total expectation.

For the subset of SNPs under parallel selection, we suppose cor(*X*_1_, *X*_2_) = *ρ* and for the remainder of the SNPs we suppose no correlation, or cor(*Y*_1_, *Y*_2_) = 0. Then the genome-wide correlation is cor(*Z*_1_, *Z*_2_) = *p ρ*. Here, *p* is the proportion of SNPs under parallel selection and it can be estimated as *p* = cor(*Z*_1_, *Z*_2_) ÷ *ρ*. We do not know what the correlation *ρ* between clinal and seasonal change should be for the variants under selection, but since it is expected to be –1 < *ρ* < 0, estimating *p* as −cor(*Z*_1_, *Z*_2_) is conservative (also see Fig. S4). Recall we assume *ρ* is negative because the climate becomes colder in higher latitudes, but it gets warmer from spring to fall.

Note our model has a few assumptions: (i) our measures of clinal and seasonal change have mean zero and variance one, which is met given we are dealing with the z-normalized regression coefficients and (ii) all polymorphisms are independent from one another. Accounting for linkage disequilibrium is notoriously complicated in genomic analyses, especially because we cannot accurately measure LD from pooled sequencing (Feder et al. 2012). However, in the previous section we showed that we are able to mitigate the effects of LD on the correlation between clinal and seasonal change using a thinning approach.

We can now readily interpret our observed correlations as proportion of SNPs under parallel selection (ignoring the negative sign). Using our thinned estimates, the patterns uncovered here are consistent with 3.78% of the common, autosomal variants being under parallel selection. It is curious our estimated proportion is close to previous estimates for the proportion of clinal (3.7% in Machado et al. 2015) and seasonal SNPs (~4% in Machado et al. 2021), both of which were obtained using different signals (using regression analyses P-values, as opposed to correlations between clinal and seasonal change).

## Discussion

Clinal patterns have been observed in both phenotypic and genotypic traits in many different species (Hancock et al. 2008; Baxter et al. 2010; Adrion et al. 2015). Especially in systems in which there is collinearity between the axis of gene flow and environmental heterogeneity, disentangling the contribution of selection and demography in producing clines is not trivial. Detecting seasonal cycling in allele frequencies is also challenging, mostly because the effect size is likely to be small and the environment may change unpredictably within seasons. The environment changes similarly with latitude and through seasons, so by jointly modelling spatial and temporal changes in allele frequency it may be possible to disentangle the role of adaptive and non-adaptive processes. Here, we showed that clinal and seasonal changes are correlated across the *D. melanogaster* genome, suggesting natural selection plays an important role in structuring allele frequencies over latitude and seasons.

Demographic processes are expected to impact the genome as a whole, but the effects of selection are stronger in regions with higher densities of functional sites (Andolfatto 2005). Consistent with this expectation, we found that correlation between clinal and seasonal change varies across genomic regions, being stronger in coding regions (Fig. 2A,C). We derived a way to biologically interpret our statistic of interest, the correlation between clinal and seasonal change. We found that allele frequency changes in roughly 3.7% of common, autosomal SNPs could be driven by natural selection.

Because we expect selection to intensify linkage disequilibrium, the correlation between clinal and seasonal variation could be mostly driven by a few large effect loci. Segregating inversions are known to underlie much of climatic adaptation in *D. melanogaster* (Fabian et al. 2012, Kapun et al. 2014, 2016), therefore we investigated how much of our signal depended on inversion status. We found the correlation between clinal and seasonal change to be stronger surrounding common inversions, highlighting the role of selection in driving frequency changes in common, cosmopolitan inversions in *D. melanogaster*. The correlation between clinal and seasonal change is particularly high for SNPs near In(3R)Payne breakpoints, an inversion known to be associated to phenotypes relevant to adaptation to cooler climates (reviewed in Kapun et al. 2019). Nevertheless, clinal and seasonal change are significantly correlated for SNPs far from inversion breakpoints, suggesting loci involved in adaptation at the spatial and seasonal scales are not restricted to inversions. We also controlled for autocorrelation along chromosomes and found that the effects of linkage disequilibrium are rather strong, but they decay rapidly and seem to return to background levels after 5kb (Fig. 3A-C). Indeed, the correlation between clinal and seasonal change remains rather strong even after accounting for LD, suggesting parallel selection acts pervasively across the genome.

Population substructure and migration could be causing seasonal variation in allele frequency in *D. melanogaster*. For example, rural populations of *D. melanogaster* in temperate regions could collapse during the winter and recover from spring to fall. However, reproductive diapause cycles in orchards and reaches high frequencies early in the spring, whereas its frequency in urban fruit markets in Philadelphia is much lower (Schmidt and Conde 2006). Another possibility is that seasonal variation is produced by migration of flies from the south in the summer, and from the north in the winter. There is little evidence of long-range migration in *D. melanogaster*, though this process seems important in *D. simulans* (Bergland et al. 2014; Machado et al. 2016). *D. melanogaster* have been shown to survive and reproduce during winter season in temperate regions, so flies can withstand a harsh winter season and be subject to selection (Mitrovski and Hoffmann 2001; Hoffmann et al. 2003, Rudman et al. 2019). These seasonal patterns have been replicated in many populations across North America and Europe (Machado et al. 2021), bolstering the argument for seasonal adaptation. Given the patterns we uncovered here are the result of subtle, but repeatable changes across multiple seasons, it is hard to imagine that selection is not the main causing force, even if it is acting to maintain cryptic population structure within each location.

Differential admixture from Europe and Africa to the ends of the clines cannot plausibly explain the parallel clinal and repeatably seasonal changes in allele frequencies (Duchen et al. 2013; Kao et al. 2015; Bergland et al. 2016), because variation over seasonal time scales is less affected by broader scale migration patterns. Further, the evidence for secondary contact in Australia is quite weak (see Bergland et al. 2016), but we show that clinal variation in Australia is correlated with seasonal variation in Pennsylvania (Fig. S3). Secondary contact may have contributed ancestral variation, which has since been selectively sorted along the cline (Flatt 2016). Consistent with this interpretation that selection mediates admixture in *D. melanogaster*, it has been found that the proportion of African ancestry is lower in low recombination regions (Pool 2015).

An important mechanism that can cause clinal patterns has been neglected from recent discussions of clinal variation in *Drosophila*. The latitudinal variation on the onset of seasons can produce clines, a phenomenon termed “seasonal phase clines” (Roff 1980; Rhomberg and Singh 1986). Under this model, a correlation between clinal and seasonal change is expected. Our latitudinal samples were all collected within one month of difference (during the spring), and so our observations could be partially explained by differences in the seasonal phase. Our data does not allow for proper disentangling of seasonal phase clines from parallel environmental change but change on the onset of seasons alone cannot explain our results. We found that latitude is usually a much better predictor of allele frequency differences (Fig. 1A-B), and the magnitude of change along the cline is much greater than what we found within a population across seasons (9.2% vs. 2.64%). We show that including Maine samples (which were obtained in the fall, in contrast to all other samples that were sampled in the spring) in our analyses does not meaningfully change our main results (Fig. S3). However, the rate of change (per degree of latitude) in allele frequency from Florida to Maine is smaller than the rate from Florida to Massachusetts. This could be due to differences in sampling year, but we also found the rate of change in frequency between Virginia and Maine to be smaller than the rate between Virginia and Massachusetts (FL and ME were sampled in 2009, whereas VA and MA were sampled in 2012; Fig. S5). We believe these differences are due to the shift in seasonal phase, as the samples from Maine were collected in the fall, but all other samples were collected in the spring.

A recent study suggested the temporal changes in allele frequency reported in Bergland et al. (2014) is only weakly consistent with seasonal selection (Buffalo et al. 2019). Consecutive spring-fall pairs showed some signal of adaptation, but the effect was small and disappeared at larger timescales (same season but across different years). Similarly, Machado et al. 2021 found that when they flipped the season labels of some samples the seasonal model fit was greatly improved. Consistent with these observations, we found there is strong temporal structure across years (Fig. 1C) and the matrix of pairwise F_ST_ shows some strong and seemingly haphazard temporal events (e.g., consider the entries for PA_07_2010 and PA_07_2015 in Fig. S5). Here with an expanded seasonal set of samples, we show that the distribution of seasonal P-values is only slightly enriched for low P-values (Fig. 1B), but top seasonal SNPs are enriched for functional genic classes, when compared to datasets in which the season labels were permuted (Fig. 2B). These results highlight how difficult it is to find truly seasonal SNPs with current datasets. Once more comprehensive time series data is available, environmental heterogeneity could be explored without the need for an imperfect proxy (such as seasons).

Many species occur along spatially structured environments and show clinal variation in traits, so a question that remains open is: what is the role of selection in producing and maintaining these patterns? Seasonal variation is also ubiquitous, especially in temperate environments, so seasonal change could be an important feature of organisms that have multiple generations each year (Behrman et al. 2015). Here, we demonstrate that by integrating clinal and seasonal variation, we can discern the contributions of selection in driving allele frequency changes with the environment. Our empirical work suggests a considerable fraction of variants distributed across the genome underlie adaptation to environmental changes over space and time in *D. melanogaster*. Importantly, our findings, together with previous studies on seasonal adaptation in flies, are bound to challenge new theoretical developments on the mechanisms that are compatible with rapid and polygenic responses to changes in the environment (e.g., Wittmann et al. 2017).

## Author contributions

M.F.R. and R.C. designed the research. M.F.R. performed the research and analyzed the data. M.F.R., M.D.V. and R.C. discussed the results and conclusions. M.F.R. wrote the manuscript with input from M.D.V and R.C.

## Acknowledgements

We would like to thank all the members of the Cogni Lab for input and support throughout the development of this work. We thank Diogo Meyer, Paul Schmidt, Reinaldo Azevedo and Peter Ralph for discussions and three anonymous reviewers for comments on the manuscript. Funding for this work was provided by São Paulo Research Foundation (FAPESP) (13/25991-0 and 17/02206-6 to RC, 15/20844-4 to MDV, 16/01354-9 and 17/06374-0 to MFR), and CNPq (307015/2015-7 and 307447/2018-9 to RC), and a Newton Advanced Fellowship from the Royal Society to RC.

## Data accessibility

All the data used in this project are available on the NCBI Short Read Archive (BioProject accession numbers PRJNA256231 and PRJNA308584, and NCBI Sequence Read Archive SRA012285.16).

## Supplementary information

**Figure S1.**
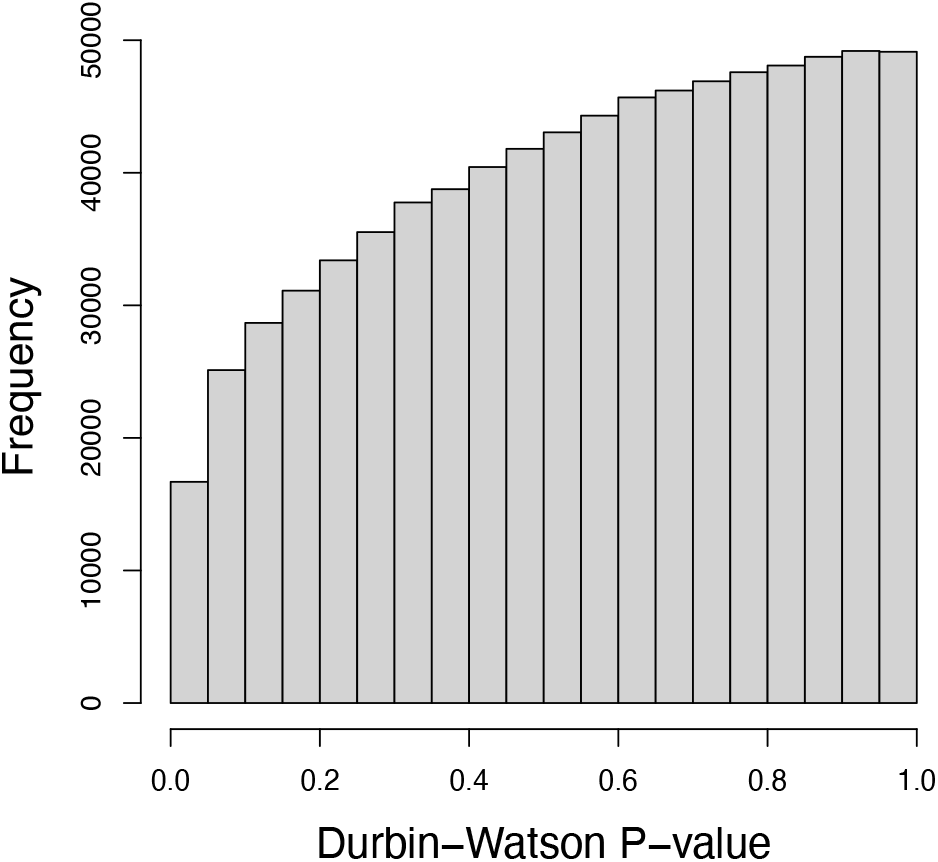
Distribution of Durbin-Watson P-values. We tested whether there is autocorrelation in the residuals of the seasonal generalized linear models.

**Figure S2.**
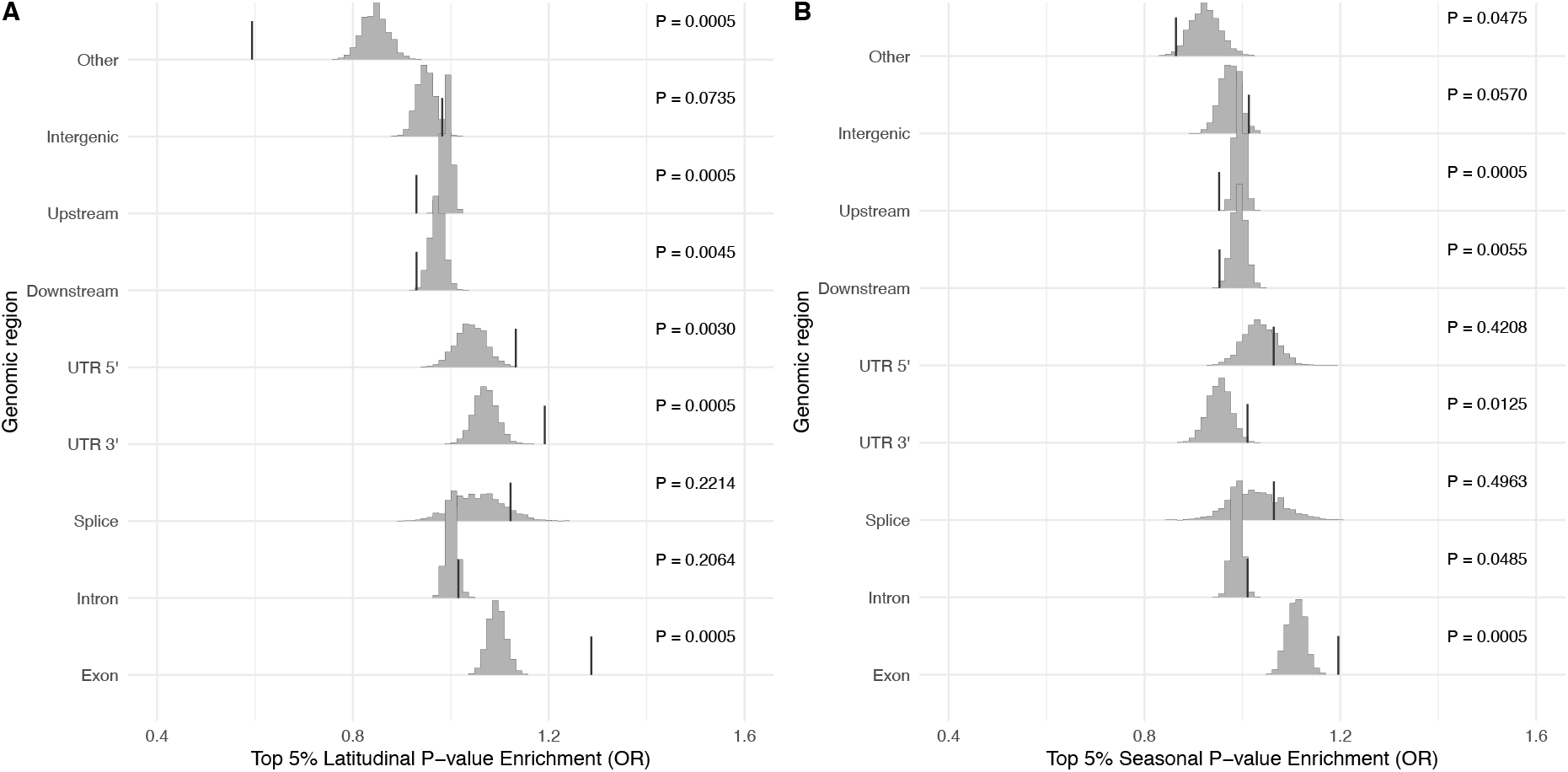
Enrichment of top 5% SNPs in each genic class for A) latitudinal P-value, B) seasonal P-value. Histograms show the distribution of odds ratios when season labels were permuted, and the vertical bars indicated the observed odds ratio.

**Figure S3.**
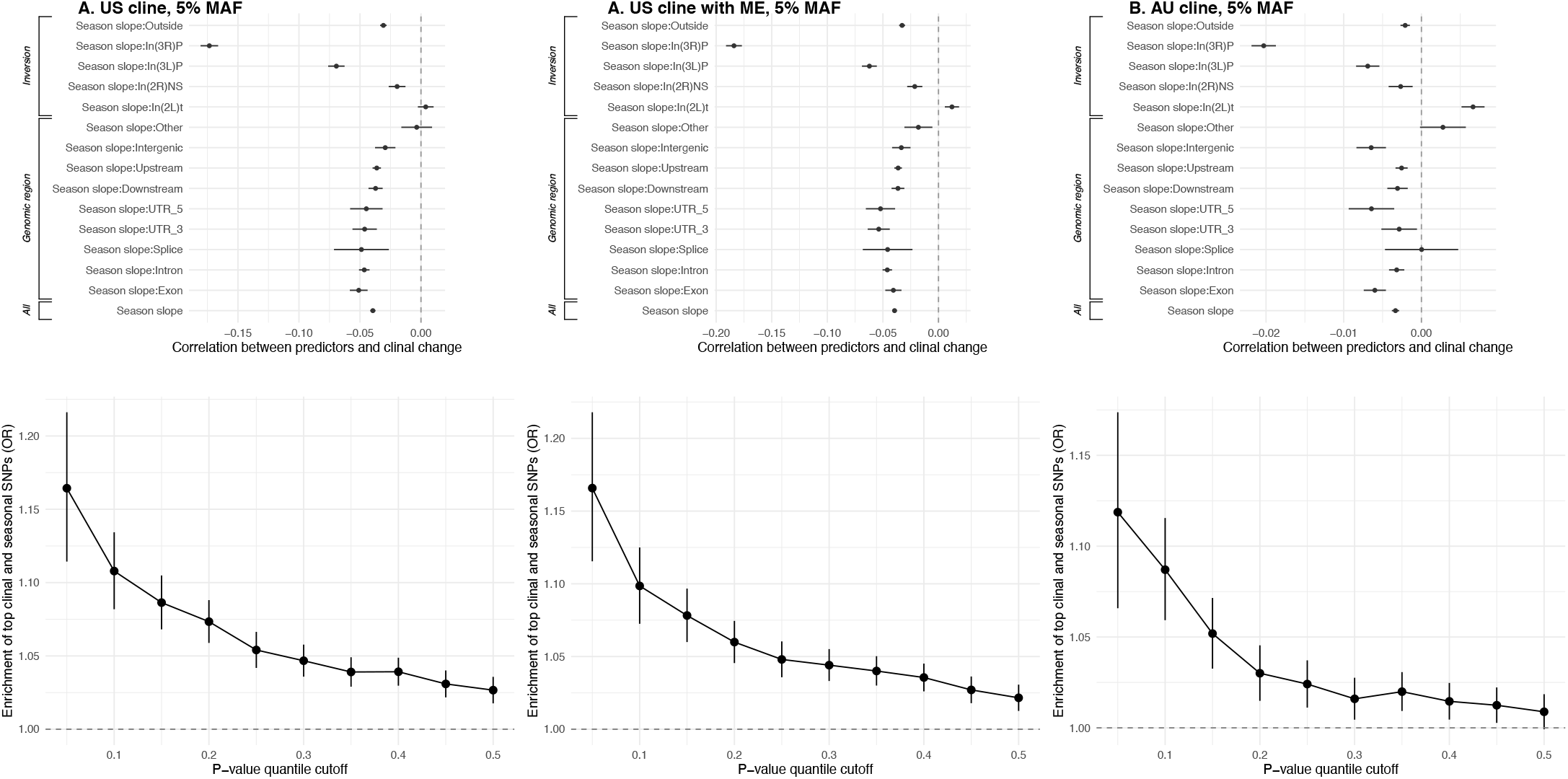
Correlation between clinal and seasonal change using A) 5% allele frequency cutoff (main results), B) latitudinal samples from Maine, which were collected in the summer and C) samples from Australia to calculate latitudinal change.

**Figure S4.**
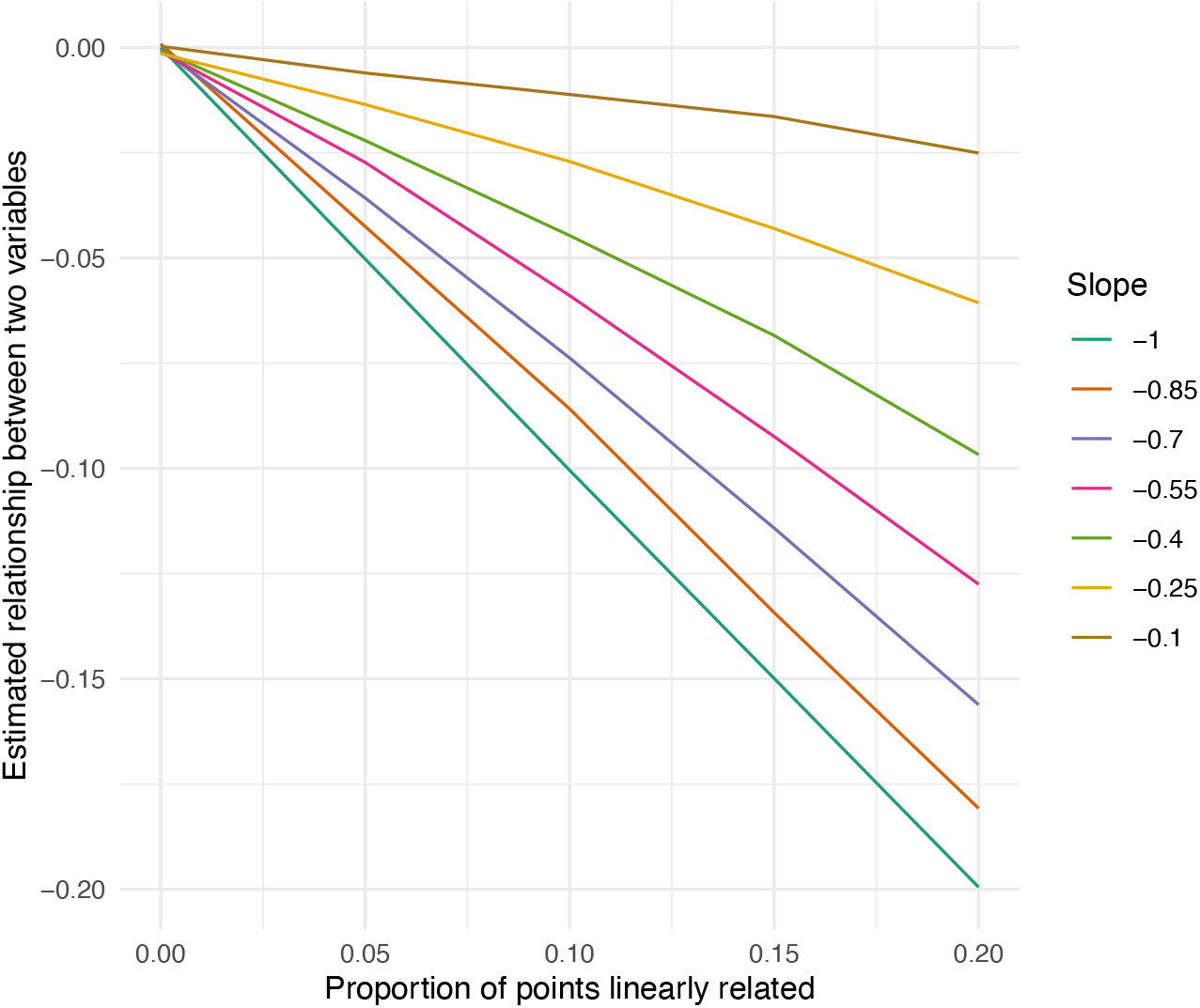
Relationship between proportion of points linearly related (in comparison to points that are not related) and the correlation between two z-normalized variables. Each color represents a different assumed degree of linear relatedness (ranging from −1 to −0.1). Note how for a given estimated relationship between two variables, the slope we assume (−1) results in the smallest possible proportion of points linearly related, demonstrating that our test is conservative.

**Figure S5.**
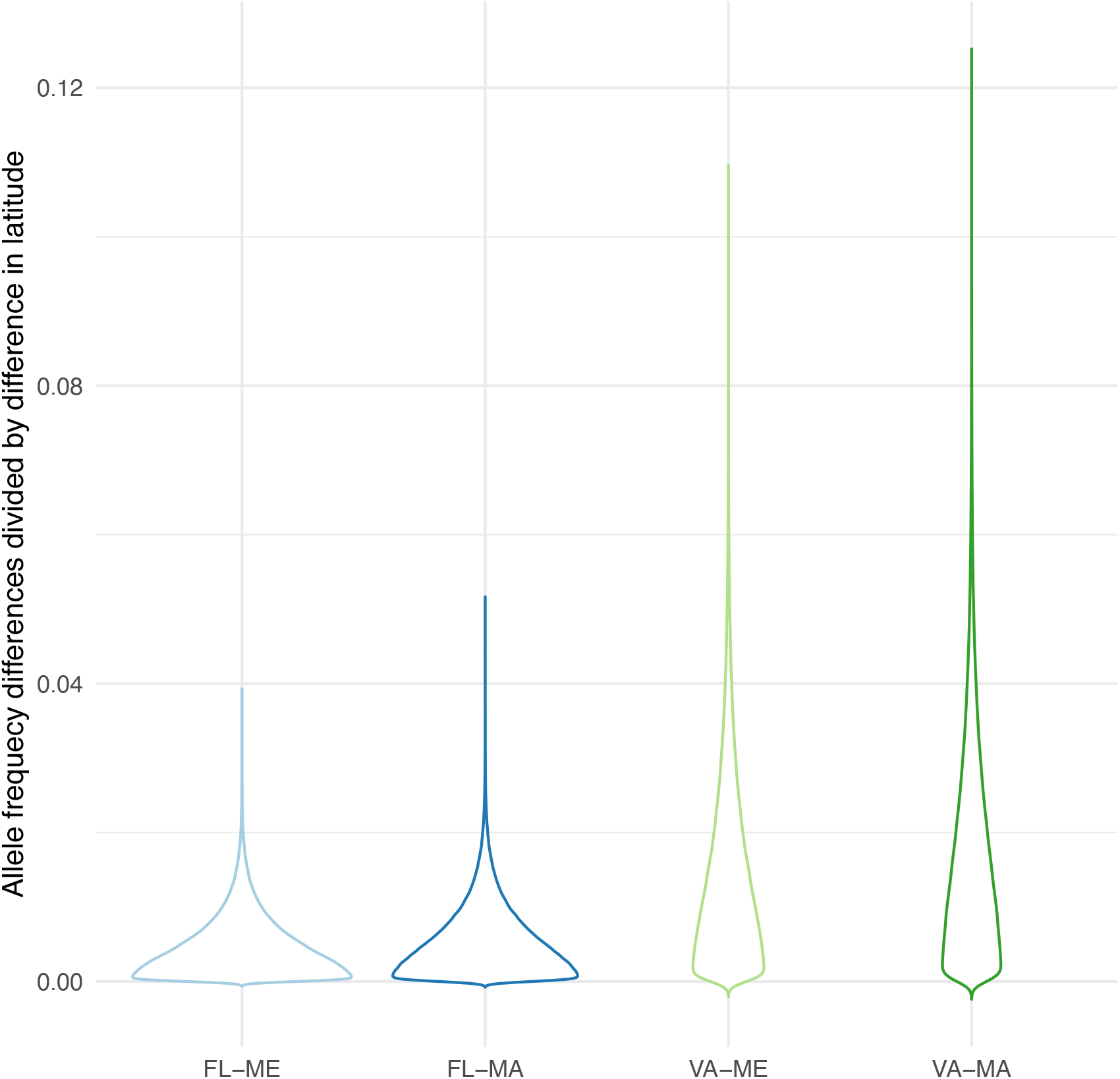
Rate of allele frequency differences between two populations (normalized by difference in latitude). Note the rates are smaller for comparisons involving the samples from Maine. FL: Florida (July 2008 and 2010), ME: Maine (October 2009), MA: Massachusetts (July 2012), VA: Virginia (July 2012).

**Figure S6.**
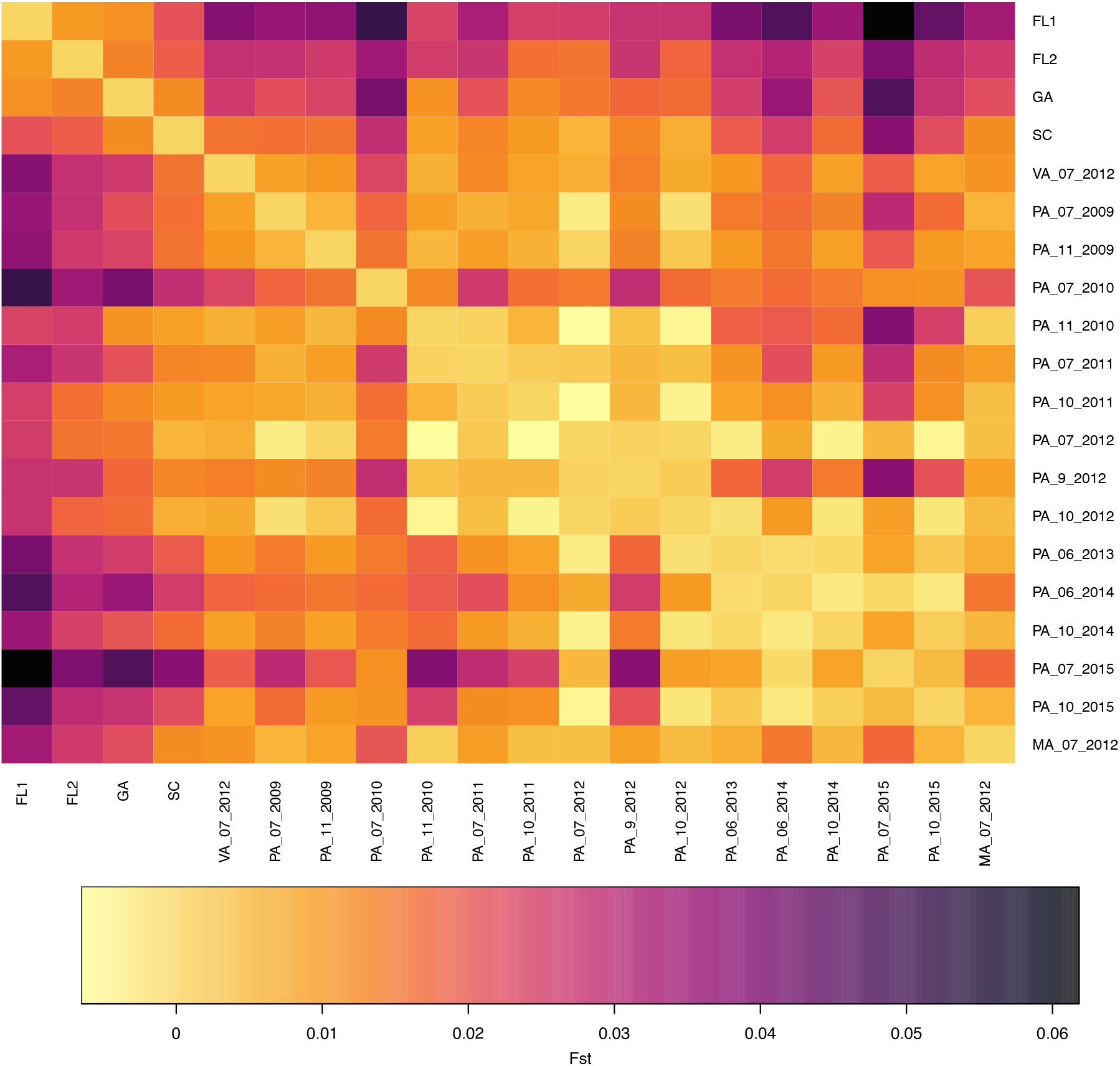
Mean pairwise F_ST_ values for all US samples included in our main analyses. Refer to Table S1 for more information on each population.

**Table S1.** Information of the samples used in this study.

| Access<br>ion # | Sample<br>name | Location | Latit<br>ude | Collect<br>ion<br>date | #<br>Fli<br>es | Medi<br>an<br>depth | Mo<br>nth | Seas<br>on | Seas<br>onal<br>set | Clin<br>al<br>set |
| --- | --- | --- | --- | --- | --- | --- | --- | --- | --- | --- |
| SRR11<br>77951 | AU_LO1 | Queensland,<br>Australia | 16.90 | 2004 | 17 | 10 |  |  |  |  |
| SRR11<br>77952 | AU_LO2 | Queensland,<br>Australia | 16.90 | 2004 | 17 | 12 |  |  |  |  |
| SRR11<br>77953 | AU_HI1 | Tasmania,<br>Australia | 42.77 | 2004 | 15 | 10 |  |  |  |  |
| SRR11<br>77955 | AU_HI2 | Tasmania,<br>Australia | 42.77 | 2004 | 15 | 14 |  |  |  |  |
| SRR15<br>25685 | FL1 | Homestead,<br>FL | 25.47 | Jul-08 | 39 | 59 | 7 | Spri<br>ng | 0 | 1 |
| SRR15<br>25694 | FL2 | Homestead,<br>FL | 25.47 | Jul-10 | 48 | 37 | 7 | Spri<br>ng | 0 | 1 |
| SRR15<br>25698 | ME1 | Bowdoinha<br>m, ME | 44.02 | Oct-09 | 75 | 86 | 9 | Fall | 0 | 0 |
| SRR20<br>06283 | ME2 | Bowdoinha<br>m, ME | 44.02 | Oct-09 | 75 | 22 | 9 | Fall | 0 | 0 |
| SRR15<br>25695 | GA | Hahira, GA | 30.99 | Jul-10 | 51 | 101 | 7 | Spri<br>ng | 0 | 1 |
| SRR15<br>25696 | SC | Euatwville,<br>SC | 33.40 | Jul-10 | 48 | 83 | 7 | Spri<br>ng | 0 | 1 |
| SRR35<br>90551 | VA_07_2<br>012 | Charlottesv<br>ille, VA | 38.03 | Jul-12 | 69 | 70 | 7 | Spri<br>ng | 0 | 1 |
| SRR39<br>39095 | PA_06_2<br>013 | Linvilla, PA | 39.88 | Jun-13 | 54 | 37 | 6 | Spri<br>ng | 0 | 1 |
| SRR35<br>90557 | MA_07_<br>2012 | Lancaster,<br>MA | 42.46 | Jul-12 | 90 | 51 | 7 | Spri<br>ng | 0 | 1 |
| SRR15<br>25768 | PA_07_2<br>009 | Linvilla, PA | 39.53 | Jul-09 | 55 | 186 | 7 | Spri<br>ng | 1 | 0 |
| SRR15<br>25769 | PA_11_2<br>009 | Linvilla, PA | 39.53 | Nov-09 | 74 | 66 | 11 | Fall | 1 | 0 |
| SRR15<br>25770 | PA_07_2<br>010 | Linvilla, PA | 39.53 | Jul-10 | 11<br>6 | 17 | 7 | Spri<br>ng | 1 | 0 |
| SRR15<br>25771 | PA_11_2<br>010 | Linvilla, PA | 39.53 | Nov-10 | 33 | 76 | 11 | Fall | 1 | 0 |
| SRR15<br>25772 | PA_07_2<br>011 | Linvilla, PA | 39.53 | Jul-11 | 75 | 53 | 7 | Spri<br>ng | 1 | 0 |
| SRR15<br>25773 | PA_10_2<br>011 | Linvilla, PA | 39.53 | Oct-10 | 47 | 74 | 10 | Fall | 1 | 0 |
| SRR35<br>90560 | PA_10_2<br>012 | Linvilla, PA | 39.53 | Oct-12 | 10<br>0 | 25 | 10 | Fall | 1 | 0 |
| SRR35<br>90561 | PA_07_2<br>012 | Linvilla, PA | 39.53 | Jul-12 | 11<br>5 | 59 | 7 | Spri<br>ng | 1 | 0 |
| SRR35<br>90563 | PA_9_20<br>12 | Linvilla, PA | 39.53 | Sep-12 | 50 | 55 | 9 | Fall | 1 | 0 |
| SRR39<br>39096 | PA_10_2<br>014 | Linvilla, PA | 39.88 | Oct-14 | 50 | 103 | 10 | Fall | 1 | 0 |
| SRR39<br>39097 | PA_06_2<br>014 | Linvilla, PA | 39.88 | Jun-14 | 92 | 68 | 6 | Spring | 1 | 0 |
| SRR39<br>39098 | PA_10_2<br>015 | Linvilla, PA | 39.88 | Oct-15 | 52 | 97 | 10 | Fall | 1 | 0 |
| SRR39<br>39099 | PA_07_2<br>015 | Linvilla, PA | 39.88 | Jul-15 | 74 | 215 | 7 | Spring | 1 | 0 |

**Table S2.** Summary of a regression of z-normalized latitude regression coefficients against z-normalized season regression coefficients genome-wide. CI stands for 95% confidence interval.

| Latitudinal slope |  |  |  |
| --- | --- | --- | --- |
| *Predictors* | *Estimates* | *CI* | *p* |
| Intercept | 0.014 | 0.012 – 0.016 | **<0.001** |
| Seasonal slope | -0.039 | -0.042 – -0.037 | **<0.001** |
| Observations | 798196 |  |  |

**Table S3.** Summary of a regression of z-normalized latitude regression coefficients against z-normalized season regression coefficients for each genic class. CI stands for 95% confidence interval.

| Latitudinal slope |  |  |  |
| --- | --- | --- | --- |
| *Predictors* | *Estimates* | *CI* | *p* |
| Intercept | 0.014 | 0.011 – 0.016 | **<0.001** |
| Seasonal slope: Intergenic | -0.029 | -0.038 – -0.021 | **<0.001** |
| Seasonal slope: Exon | -0.051 | -0.058 – -0.044 | **<0.001** |
| Seasonal slope: Intron | -0.046 | -0.051 – -0.042 | **<0.001** |
| Seasonal slope: Downstream | -0.037 | -0.043 – -0.031 | **<0.001** |
| Seasonal slope: Upstream | -0.036 | -0.040 – -0.033 | **<0.001** |
| Seasonal slope: UTR_3 | -0.046 | -0.056 – -0.036 | **<0.001** |
| Seasonal slope: UTR_5 | -0.045 | -0.058 – -0.032 | **<0.001** |
| Seasonal slope:Splice | -0.049 | -0.071 – -0.026 | **<0.001** |
| Seasonal slope:Other | -0.004 | -0.016 – 0.009 | 0.584 |
| Observations | 798196 |  |  |

**Table S4.** Summary of a regression of z-normalized latitude regression coefficients against z-normalized season regression coefficients for each SNPs surrounding inversion breakpoints. CI stands for 95% confidence interval.

| *Predictors* | **Latitudinal slope** |  |  |
| --- | --- | --- | --- |
|  | *Estimates* | *CI* | *p* |
| Intercept | 0.014 | 0.012 – 0.016 | **<0.001** |
| Seasonal slope:In(2L)t | 0.004 | -0.003 – 0.010 | 0.235 |
| Seasonal slope:In(2R)NS | -0.020 | -0.026 – -0.013 | **<0.001** |
| Seasonal slope:In(3L)P | -0.069 | -0.076 – -0.063 | **<0.001** |
| Seasonal slope:In(3R)P | -0.173 | -0.181 – -0.166 | **<0.001** |
| Seasonal slope:Outside | -0.031 | -0.033 – -0.028 | **<0.001** |
| Observations | 798196 |  |  |

